# A glimpse into the diverse cellular immunity against SARS-CoV-2

**DOI:** 10.1101/2021.02.17.431750

**Authors:** Cheng-Wei Chang, Yuchen Liu, Cheng Jiao, Hongwei Liu, Xiaochuan Chen, Lung-Ji Chang

**Affiliations:** Shenzhen Geno-Immune Medical Institute, Shenzhen, China; University of Electronic Science and Technology of China, Sichuan, China; Wellness Medical Center, New Jersey, USA

**Author notes:** Correspondence*: Lung-Ji Chang, Ph.D., Shenzhen Geno-Immune Medical Institute, 6 Yuexing 2nd Rd., 2^nd^ Floor, Nanshan Dist. Shenzhen, Guangdong Province, China 518057.

**Keywords:** Covid-19, SARS-CoV-2, Cellular Immunity, Vaccine

## Abstract

Severe acute respiratory syndrome coronavirus 2 (SARS-CoV-2)-specific cellular immune response may prove to be essential for long-term immune protection against the novel coronavirus disease 2019 (COVID-19). To assess COVID-19-specific immunity in the population, we synthesized selected peptide pools of SARS-CoV-2 structural and functional proteins, including Spike (S), Membrane (M), Envelope (E), Nucleocapsid (N) and Protease (P) as target antigens. Survey of the T cell precursur frequencies in healthy individuals specific to these viral antigens demonstrated a diverse cellular immunity, including high, medium, low and no responders. This was further confirmed by *in vitro* induction of anti-SARS-CoV-2 T cell immune responses using dendritic cell (DC)/T cell coculture, which supported the corresponding T cell precursor frequencies in each of the individuals tested. In general, the combination of all five viral antigen pools induced the strongest cellular immune response, yet individual donors responded differently to different viral antigens. Importantly, *in vitro* restimulation of the T cells with the DC-peptides induced increased anti-viral immune responses in all individuals even in the no responders, suggesting that repeated antigen stimulation could elicit a broad protection in immune naïve population. Our analysis recapitulates the critical role of cellular immunity in fighting COVID-19 and the importance of analyzing anti-SARS-CoV-2 T cell response in addition to antibody response in the population.

**Importance:** Facing the rapid evolving SARS-CoV-2 variants in the world, current emphasis on antibody-producing vaccines needs a quick revisit. The virus-specific cellular immunity may prove to be essential for long-term protection against COVID-19. This study designed a series of antigenic peptides encompassing the conserved and/or essential domains of Spike (S), Membrane (M), envelope (E), Nucleocapsid (N) and Protease (P) as targets to assess Covid-19-specific immunity in the population. The results demonstrated a diverse cellular immunity, including high, medium, low and no responders. This was verified by *in vitro* generation of anti-SARS-CoV-2 T-cells from these subjects. The study suggested that individuals responded differently to the different viral antigens, and importantly, repeated stimulation could produce virus specific T cells in all individuals, including the no responders. This study illustrates the needs for assessing anti-viral cellular immunity in addition to antibody response in the general population.

## Introduction

The main clinical manifestations of COVID-19 are fever, fatigue, dry cough and other respiratory and systemic symptoms. Recent surveys show that most of the virus carriers (more than 60%) have no symptoms and do not even realize that they are infected with the virus. About 25% of the carriers have very mild symptoms and can self-heal. Only about 15% of the infected people develop serious condition with body temperature over 38°C, mostly in adults over 65 and those with certain preexisting conditions such as hypertension, diabetes, and obesity.

Serious complications of Covid-19 include acute respiratory distress syndrome (ARDS), acute heart injury and secondary infections (1, 2). The SARS-CoV-2 virus can stimulate the innate immune system of patients, resulting in the release of cytokines and immune effectors locally and systemically, causing cytokine storm and acute inflammatory response (3). This can lead to increased systemic vascular vulnerability, causing acute respiratory distress and multiple organ failure (4, 5). Lymphocytopenia is a common feature in patients with severe COVID-19, accompanied by a sharp decrease in the number of CD4 and CD8 T cells, B cells, and natural killer (NK) cells (6, 7), indicating that the pathogenesis of the new SARS-CoV-2 virus is highly correlated with host immunity, which may impact the development of anti-viral therapies and vaccines.

Antibody-dependent enhancement (ADE) of viral entry has been a major concern for viral epidemiology and vaccine development. Most of the conventional vaccines are aimed at establishing humoral immunity, with the spike protein of the SARS-CoV-2 virus as the main target. There has been concerns for the ADE effect with the COVID-19 vaccines, where antibodies may facilitate viral entry into host cells and enhance viral infection, as has been observed in the HIV-1 vaccine development (8, 9). Furthermore, the humoral immunity may coexist with viremia for a prolonged period and ineffective in virus clearance, although sera from COVID-19 survivors were reported to be capable of clearing the virus in most of the recipient patients (10, 11). Recent studies reveal the existence of broad virus-specific T cell responses in asymptomatic carriers, which may highlight a critical role of cellular immunity in development of the COVID-19 vaccines (12, 13).

To control viral infection, the cytotoxic T lymphocytes (CTLs) reactive to specific viral antigens has proven to be an essential contributor. *Ex vivo* expanded antigen-specific T cells targeting cytomegalovirus (CMV), Epstein-Barr virus (EBV) and adenovirus have been successfully applied to treating hematopoietic stem cell or solid organ transplant patients who have developed post-transplant viral diseases (14, 15). The immunogenic components of a virus include a complex number of epitopes derived from the multiple viral proteins. Here we tested the immunogenicity of the various SARS-CoV-2 structural and functional proteins using pooled peptides from the invididual proteins to evaluate the virus-specific cellular immunity in healthy volunteers. The results illustrated a glimpse into the diverse host immunity in the population including no, low, medium and high responders to the different SARS-CoV-2 proteins. Extended *in vitro* immune activations using pooled viral peptides demonstrated the feasibility of eliciting an anti-viral cellular immunity even in the low and no responders.

## Results

### Synthesis of antigenic peptide pools of SARS-CoV-2 structural and functional proteins

Detailed analyses of the viral genome sequences of SARS-CoV-2 virus with the genomes of SARS virus and MERS virus revealed that the structural proteins Spike (S) and Membrane (M) of the coronavirus have high mutation rate, while the Envelope (E), Nucleocapsid (N) and Protease (P) regions are highly conserved. To identify potential vaccine targets, we analyzed the genes of all structural proteins of the virus, including S, M, E, N, and the polyprotein cleavage protease (P), and synthesized selected pools of pentadecamer peptides spanning across important functional domains of these polyproteins, including the Receptor Binding Domain (RBD) of S protein, the full-length of E membrane protein, the entire M protein as it is the most abundant protein of coronavirus, and the nucleic acid binding domain (NBD) plus the serine-arginine rich (SR) domains including the helix-turn-helix motif of N protein, and the domain III of Mpro (Figure 1 a and b). The rationale for the design of these pentadeca-peptides to encompass the selected conserved functional domains in the SARS-CoV-2 sequence is that these regions are less likely to mutate due to their functional importance, and the pentadeca-peptides can induce both class I and class II MHC-restricted T cell responses based on past experiences.

**Fig. 1.**
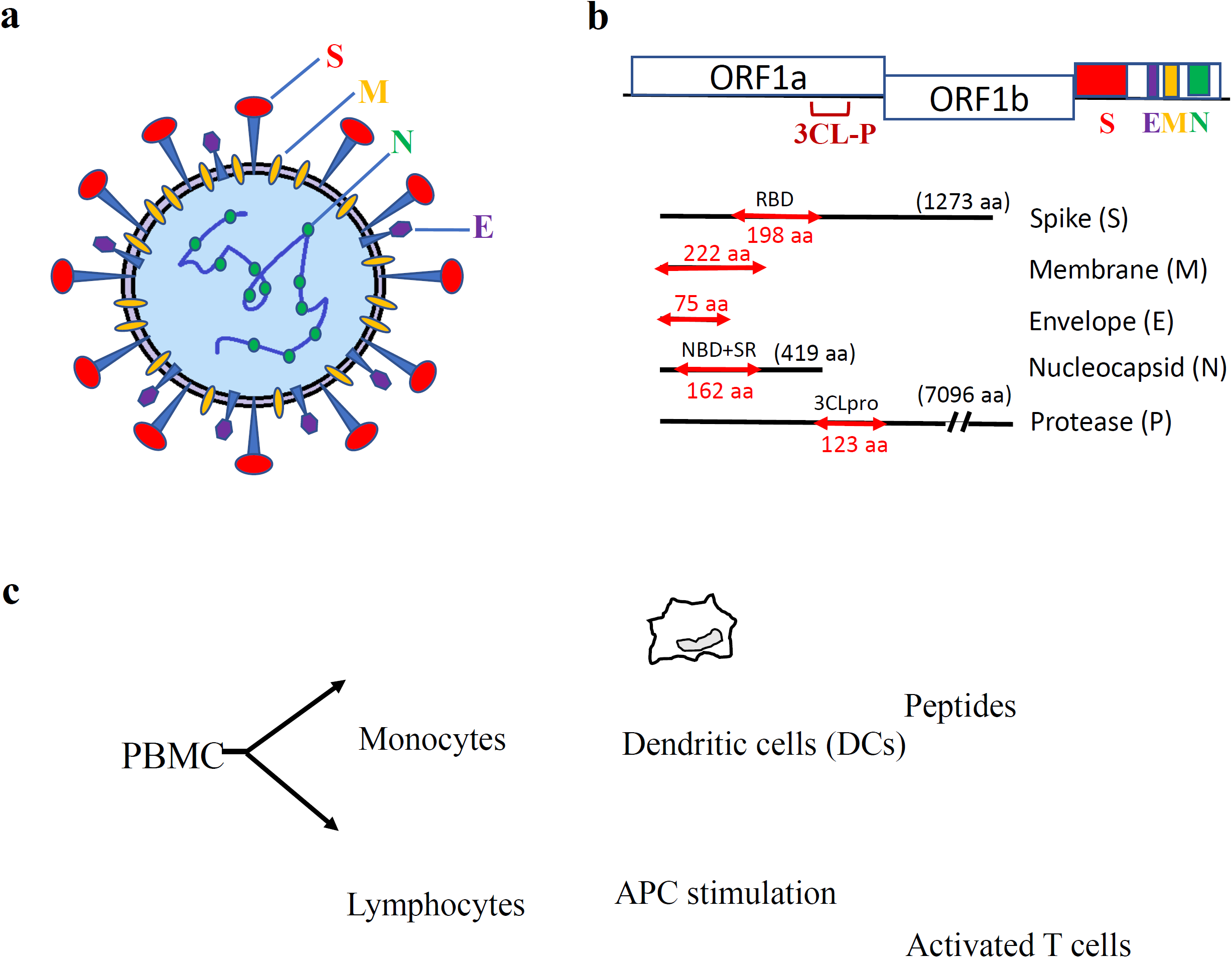
Synthetic peptides of SARS-CoV-2 S, E, M, N, and P proteins and *in vitro* generation of antigen-specific T cells. (**a**) Schematic illustration of SARS-CoV-2 virus particle. (**b**) Representative S, E, M, N, P protein domains. (**c**) Diagram of *in vitro* antigen-specific T cell generation. DCs and T lymphocytes were prepared from PBMCs and synthetic viral peptide-pulsed DCs were used as antigen presenting cells (APCs) to activate T cells.

Antigen-specific immune effector cells can be generated through antigen presentation by DCs. The protocol includes *in vitro* generation of DC, followed by antigen exposure and co-culture with autologous lymphocytes. We isolated peripheral mononuclear cells (PBMCs) from healthy volunteers, and generated matured DCs from monocytes. The specific antigen pools of SARS-CoV-2 were used to pulse DCs and then cocultured with peripheral blood lymphocyts to activate antigen-specirtic T cells as illustrated in Figure 1c.

### Assessment of SARS-CoV-2 antigen-specific precursor frequency in healthy individuals

To investigate the frequency of immune effectors to SARS-CoV-2 in healthy individuals, we examined antigen-specific T cells in 29 healthy subjects who have no known prior exposure to the SARS-CoV-2. PBMCs were isolated from the whole blood, and the pooled viral S, M, E, N and P peptides, as well as a control HIV peptide pool, were added to the PBMCs to activate immunogenic response for 17 hours, followed by IFN-γ ELISPOT analysis. The IFN-γ specific spots represented the antigen-specific T cells activated by the peptide antigens. Positive response was arbitrarily set at a 1.5-fold increase in the numbers of IFN-γ-secreting T cell spots in the test wells versus the control wells which included PBMC alone or treatment with HIV peptides.

We observed a diverse range of SARS-CoV-2-specific primary T cell frequencies in the healthy population, presented as spot forming cells in Figure 2a (Raw data in Supplementary Tables 1-3). After deducting the background value, the frequencies of antigen specific T cells in 10^5^ T cells to each peptide antigen pool for each subject are plotted in Figure 2b, and the bars represent the median value of antigen-specific T cells to each antigen pool. There was a trend that the five mixed peptide pools (SMENP) revealed the highest T cell precursor frequencies as compared with the individual viral peptide pools (Fig. 2b). The antigen specific response was also quantified based on the fold increase of spots with respect to the PBMC alone group. We set the scales as low responders if the number was 1.5-2 fold higher than the PBMC background, median responders if it was 2-3 fold higher, and high responders if it was 3 folds or higher. As shown in Figure 2c, the SMENP peptide pools induced the most robust response (24%) as compared to the other five individual peptide pools (Fig. 2c). The individuals had different response preference to the various viral antigens, with more than 60% of the tested subjects showing no detectable cellular immune response to the SARS-CoV-2 antigens (no responders). Furthermore, the high responders to the S peptide pool, mainly the RBD domain that is the preferred antigenic target used in the COVID-19 vaccine design, is only 10%, similar to those of the high responders to the M peptide pools.

**Fig. 2.**
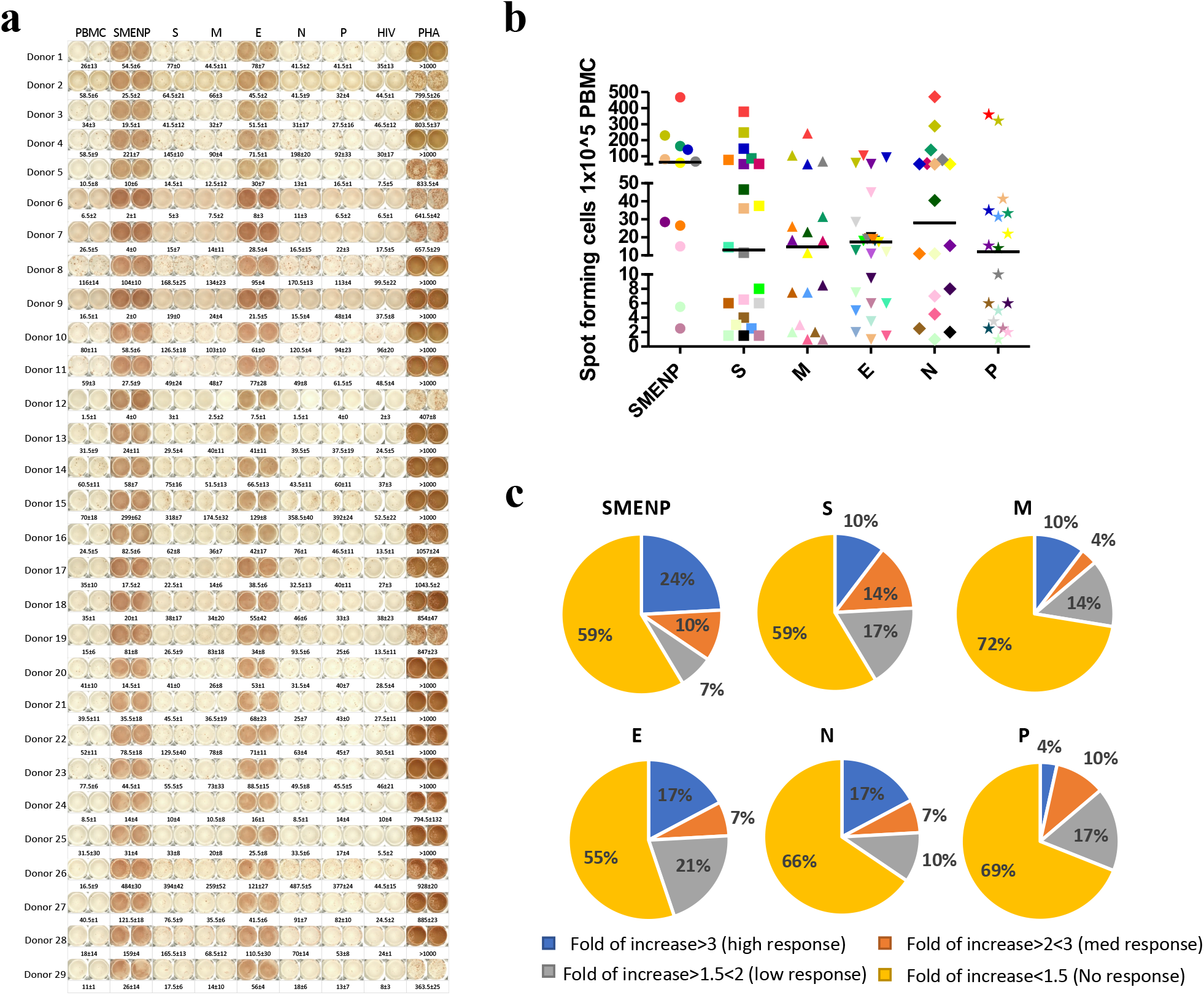
SARS-CoV-2 specific T cell precursor frequencies in healthy subjects. (**a**) IFN-γ ELISPOT assay. PBMCs of 29 healthy subjects were activated *in vitro* with individual or pooled S, E, M, N and P peptide mixes and IFN-γ reactive cells were detected. PHA activated T cells were used as positive controls. The HIV Pol peptide pool was also included as background control as all donors were HIV-negative. The group without antigen (PBMC) was also served as backgroun control. (**b**) The number of spot forming cells after background subtraction per indicated target antigens. Each symbal represents a peptide pool, and different colors represent different subjects. The bar represents median response value of all subjects to each peptide antigen pool after deducting the background value. The significance of difference P values of the SMENP peptide pool versus the individual S, M, E, N and P peptide pools are 0.2, 0.05, 0.005, 0.46, and 0.21, respectively. (**c**) Pie graph illustration of the diverse individual immune cell response to the different viral antigens based on fold of increase of spot forming cells. The color classification of individual response illustrates different ranges of IFN-γ specific spots: high responder, > 3 fold of increase in spots; medium responder, 2-3 fold of increase; low responder, 1.5-2 fold of increase; and no responder, <1.5 fold of increase.

### Activation of SARS-CoV-2-specific immune cells *in vitro*

To investigate whether T cells of different respondrers could be activated by the SARS-CoV-2 antigens, we selected donors with different responses in the precursor frequency test to perform an *in vitro* T cell activation assay. Immature DCs were generated from adherent blood monocytic cells for 5 days in the presence of GM-CSF and IL-4 (16). T cells were co-cultured with DCs pulsed with the various SARS-CoV-2 antigenic peptides, including pooled SMENP and the individual viral protein peptide pools, or a negative control HIV peptide pool, for 12 days, followed by ELISPOT analyses. The results showed that the activation potential of the individual antigen-specific T cells correlated with their corresponding T cell precursor frequencies, i.e., the high responders developed the strongest T cell response, and the no responders developed little to no response (Fig. 3a). The high responder group showed enhanced specific cellular immune response by more than 30 fold after 12 days in culture, and the medium responder group and low response group increased about 15 folds and 10 folds, respectively. On the other hand, the expansion of the antigen-specific T cells in the no responder group was relatively low (Fig. 3b). Again, there was a diversity in the individual preference in response to the various viral antigens of SARS-CoV-2, e.g, the high responder had the lowest response to the E antigen, whereas the no response donor #2 had the highest response to the E antigen.

**Fig. 3.**
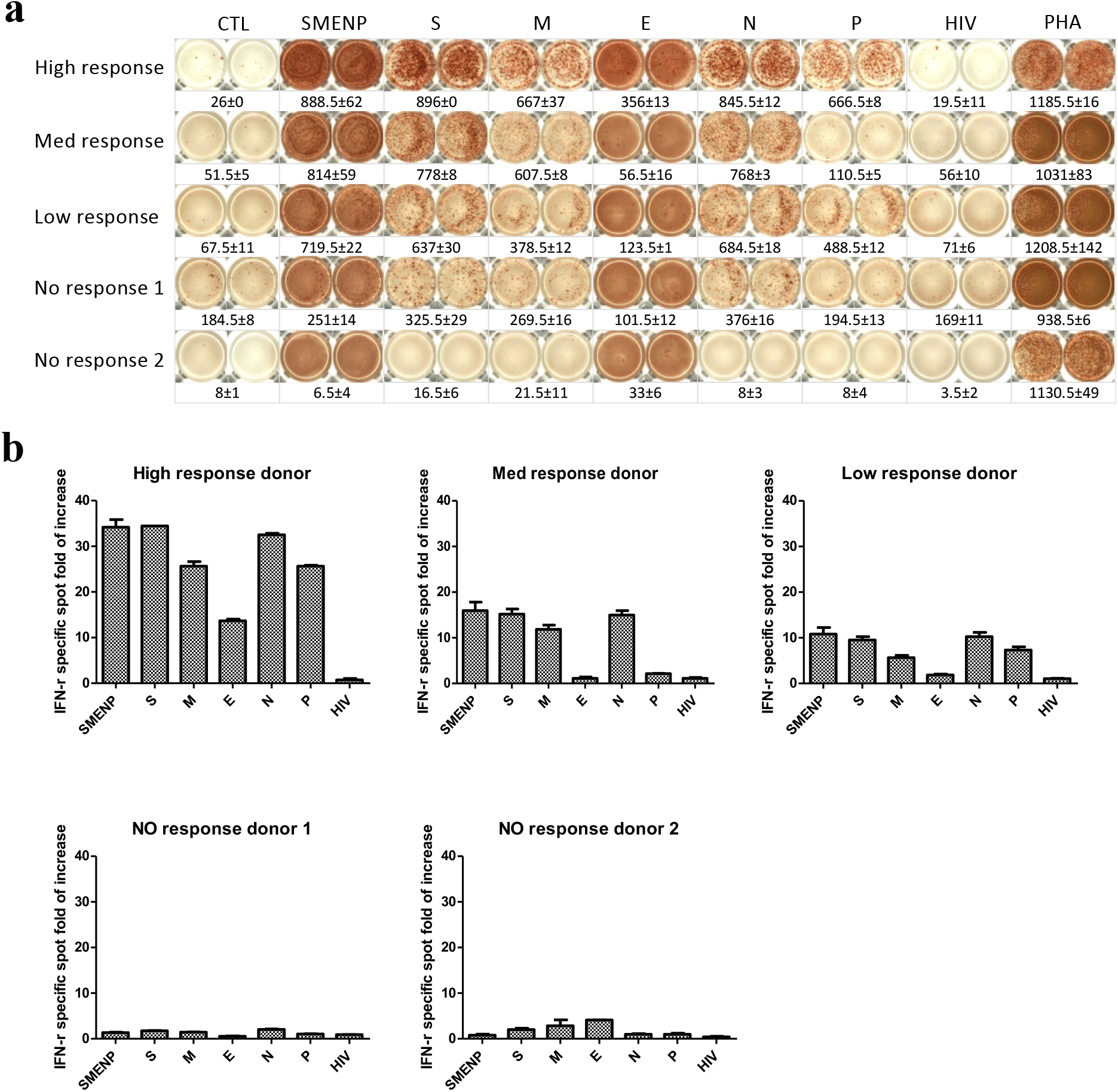
Primary activated T cells for 12 days against SARS-CoV-2 antigens. (**a**) IFN-γ ELISPOT assay of day 12 T cells from selected responders. T cells were cultured for 12 days from the different subjects including no, low, medium and high responders. PHA treatment served as positive control, and HIV peptides and CTL without antigen served as background controls. (**b**) Bar graph analysis of the IFN-γ specific T cell expansion fold of increase based on spot forming cells against the different viral antigens for the five subjects. The error bar represents the difference of the duplicate wells.

### Immune booster to enhance specific anti-viral cellular immunity

To see if the anti-viral immune response could be enhanced in the no responders by an immune booster application, we re-stimulated the *in vitro* cultured T cells from the two no responders with DCs pulsed with the same SARS-CoV-2 antigen peptide pools, and extended the culture for 30 days. ELISPOT assay was then performed to measure the specific T cell responses. As compared with the 12 day results, the background value of the non-specific cells decreased significantly (Fig. 4a), and the specific expansion of T cells increased to more than 20 folds after the secondary immune stimulation (Fig. 4b). The result suggested that individuals with low frequency of immune response to the SARS-CoV-2 virus could benefit from antigen restimulation.

**Fig. 4.**
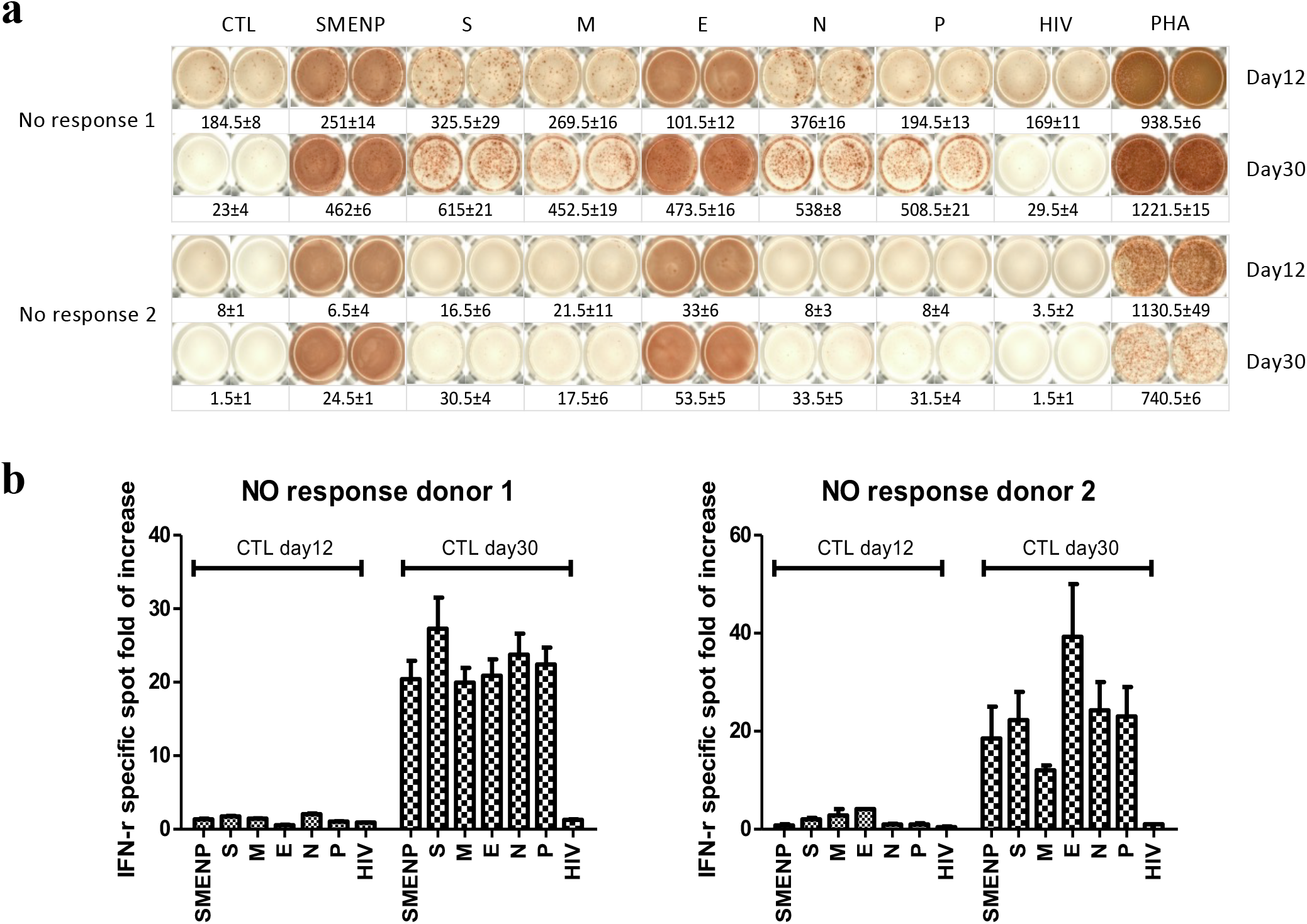
Secondary DC/T cell activation at 30 days against SARS-CoV-2 antigens. (**a**) IFN-γ ELISPOT assay of the two no responders after secondary antigen activation. The T cells after primary and secondary DC activation were cultured for 30 days from the two no response donors. PHA served as positive control, and HIV peptide pools and CTL without antigen served as background controls. (**b**) The comparison of primary day 12 versus secondary day 30 IFN-γ spot expansion folds of T cells against indicated target antigens.

### Anti-SARS-CoV-2 effector activities of the *in vitro* DC-SEMNP-activated T cells

Upon T cell receptor (TCR) engagement and stimulation by antigens in association with MHC molecules, specific immune effector functions can be demonstrated by the activation and release of specific effector molecules such as IFN-γ, TNF-α, IL-2 and CD107a (17–19). We examined the SARS-CoV-2 viral antigen-specific T cell response by intracellular staining for TNF-α, IFN-γ, IL-2 and CD107a. The generation of a SARS-CoV-2 specific T cell response was determined by comparing T cell stimulation with a control HIV peptide pool. We observed several folds of increases in the SARS-CoV-2 antigen-reactive T cells over the control T cells (Fig. 5 and supplementary Fig. 1), indicating that DCs presenting the SMENP epitopes (CTL+Covid-19) elicited a strong anti-SARS-CoV-2 T cell response in both CD4 and CD8 T cells (CD107a, IFN-γ, TNF-α and IL-2, Fig. 5b and 5c).

**Fig. 5.**
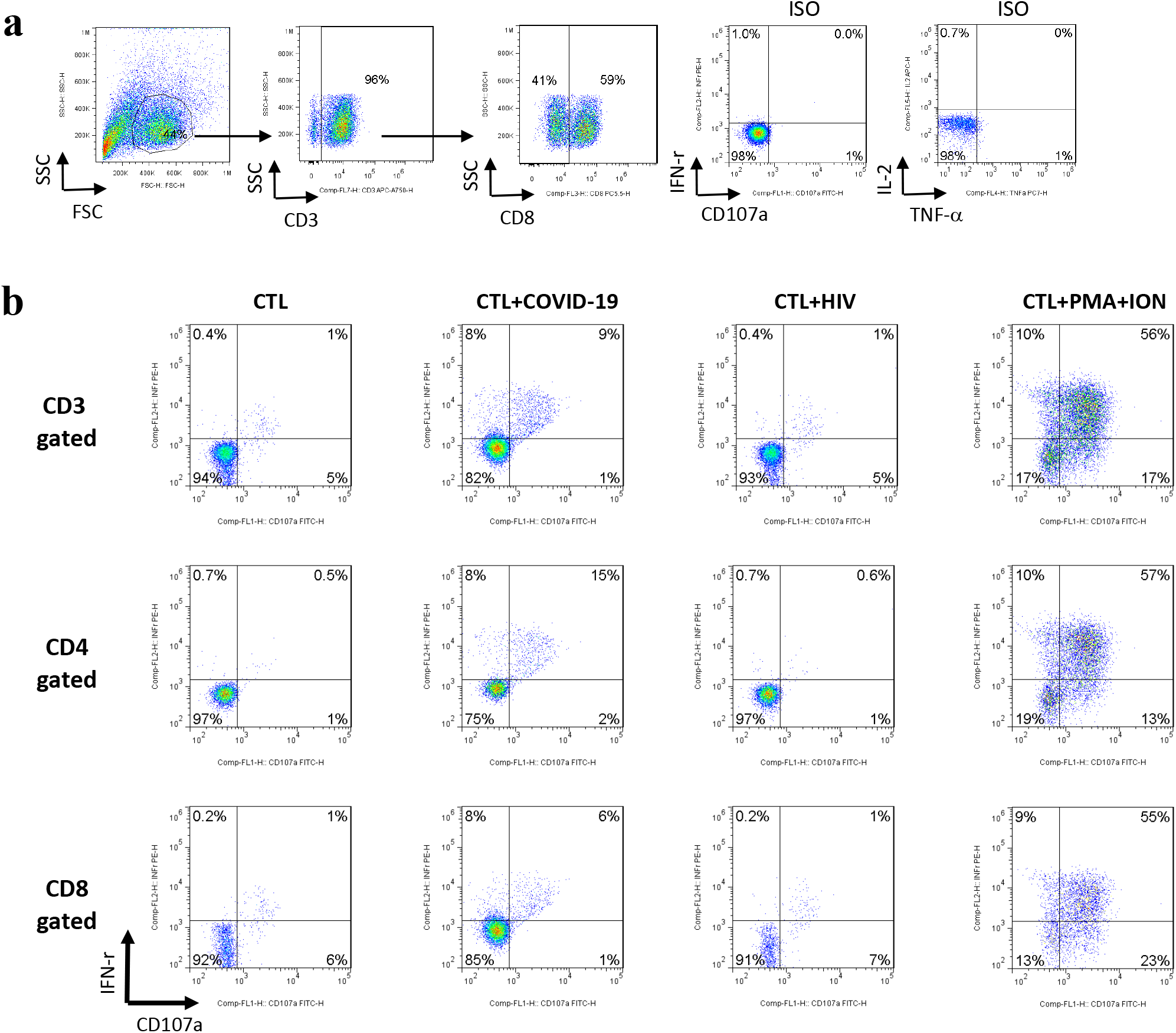
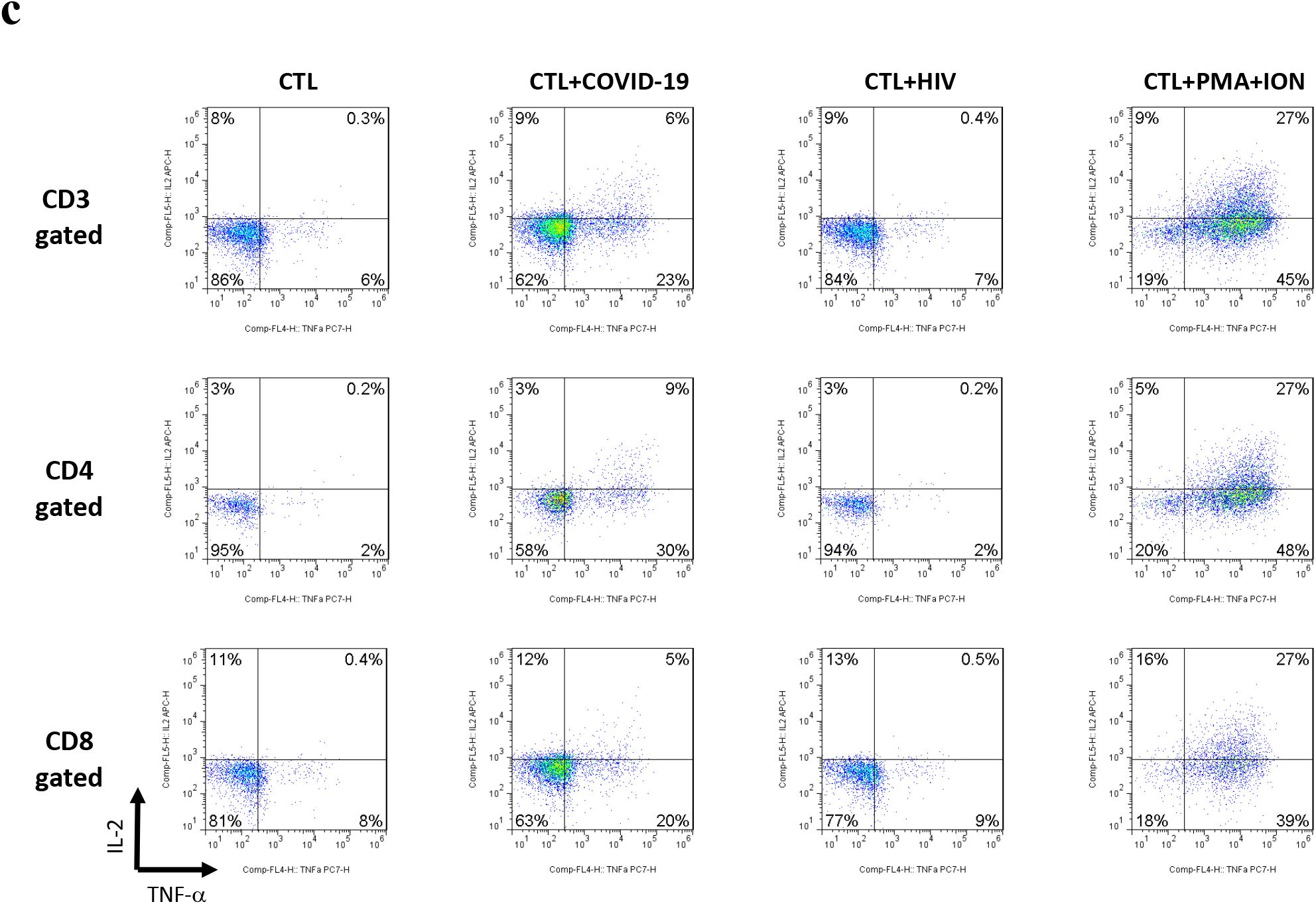
Effector function analyses of T cells against SARS-CoV-2 SEMNP antigens. After DC-T cell coculture, the viral antigen-specific CTLs from the healthy subjects were analyzed for cytokine release and CD107a degranulation by intracellular staining and flow cytometry. The TNFα, IFN-γ, IL-2 and CD107a production in T cells were illustrated in the FACS graphs after stimulations with SARS-CoV-2 antigenic peptides. PMA and ionomycin (PMA+ION) activation served as positive control, and HIV-1 peptide pools and CTL without antigen served as background controls. Flow cytometry analyses of TNFα, IL-2, IFN-γ and CD107a-positive cells in CD4-gated or CD8-gated populations of the representative subject are illustrated. (**a**) The gated CD3 and CD8 T cell populations and the control isotype antibody staining of the intracellular effectors. (**b**) Flow cytometry analysis of intracellular IFN-γ and CD107a in CD3, CD4 and CD8 gated populations. (**c**) Flow cytometry analysis of intracellular TNFα and IL-2 in CD3, CD4 and CD8 gated populations.

## Discussion

A safe and effective COVID-19 vaccine is key to end the global COVID-19 pandemic. Several vaccine candidates are currently in use, and many more are still in preclinical development (20, 21). Virus-specific memory T cells have been shown to persist for many years after infection with SARS-CoV (22, 23). Recent studies have shown that many people infected with SARS-CoV-2 virus with mild synptoms or asymptomatic status can develop T-cell-mediated immunity to the virus even without antibody response (24). In addition, ananlysis of T cell immunity to specific SARS-CoV-2 epitopes also demonstrates the existence of anti-viral T cells in un-exposed individuals (25–27). This means that the actual level of population immunity to the new coronavirus is higher than antibody positive population.

In this study, we investigated SARS-CoV-2-specific cellular immunity in healthy individuals who have no known exposure to the virus. Our results demonstrated that approximately 60-70% of tested individuals have no cellular immune response to the virus. About 20% of the tested subjects had T cell response to the virus, and a few of them had strong cellular immune response to the virus. Note that many of the subjects of this study came from Guandong province (13 of the 29), the epicenter of previous SARS epidemic. Possible prior SARS-CoV exposure cannot be excluded. It is anticipated that those with strong cellular immunity to the virus may become asymptomatic or only have mild symptoms after exposure to the virus, as well as a better prognosis, as reported by a clinical study that cellular immunity is associated with recovery from COVID-19 (12, 13). The study subjects #4, #15 and #26 had high precursor frequencies. After a retrospective survey, we discovered that subject 4, a young individual who had lived in Guangzhou, China, during the SARS-CoV peak epidemic in 2011, and this subject might have been exposed to the SARS virus at the time. The subject #15 and #26 are 65 and 39 years old, respectively, and might have been exposed to other common cold human endemic coronaviruses. The protein sequences of SARS-CoV-2 share high homology to SARS-Cov and MERS (>90%), and moderate homology to the other coronaviruses (28). The SARS-CoV-2 peptide pools designed in this study encompassed several different viral proteins, which include some highly conserved antigenic domains identical or similar to the other coronaviruses (Supplementary Table 4, CoV sequence homology comparison using BLASTP based on FASTA databases,).

It is not surprising that there is a diversity in anti-viral cellular immunity as the viral antigens are presented based on the individual HLA type. For example, it is known that HLA-B27 individuals carry an HIV-resistant phenotype (29). It is also possible that some individuals are highly suspectible to the SARS-CoV-2 infection, while others may be more resistant to the virus. Importantly, our study showed that if the immune cells were exposed to all five viral protein peptides, there was an increased overall T cell response. The design of antigenic peptides spanned across the entire protein domains which might have included regulatory T cell epitopes, and thus inhibitory T cell responses could also be induced. This study included subjects from both Southern and Northern Chinese population with high variation in HLA phenotype. While HLA-restricted immunity plays a role in the antiviral response, to further define HLA-specific SARS-CoV-2 response is out of the scope of this study. Nevertheless, extended evaluation of viral effector and regulatory antigenic domains will be helpful for anti-viral immunity studies.

The 29 subjects ranged from age 3 to 65, and 13 of them are from the Southern China Guangdong province (Supplementary Table 5). It is not clear how the age and geographic location contributed to the diverse SARS-Cov-2 cellular immunity. Interestingly, when the immune response was plotted based on the the response intensity to each viral antigens and individual age, there was a trend that the cellular immune response increased in subjects over 30 years of age (Supplementary Fig. 2). However, due to the limited subjects, the significance of this age factor awaits further investigation.

The SARS-CoV-2 virus is still evloving during the global pandemic (30), and the public immunity of different ethnics to the virus may differ. Previous studies of cellular immune protection against CMV, EBV and adenovirus have indicated that targeting at least two viral antigens to establish a wider cellular immune responses indeed can increase clinical benefits (31). Therefore, one should consider targeting multiple viral antigens, rather than a single protein such as the spike protein of SARS-CoV-2 virus, for the vaccine to have broader cellular immune responses.

A rapid cellular immune response to the SARS-CoV-2 virus may be key to the protective immunity in the host. This will ensure quick removal of the infected cells to avoid a systemic infection. It remains to be determined if a robust specific T cell response can prolong the protection against COVID-19. Nevertheless, similar scenario has been inferred from previous studies of MERS and SARS-CoV, that those individuals who have developed potent memory T cell responses after infeciton can maintain a persistent anti-viral immunity while antibody responses faded with time (32–34). It has been reported that the COVID-19 antibodies gradually decrease after infection in 90 days (35). If so, targeting cellular immunity against COVID-19 would be critical for vaccine development. Our work provides a basis for analysis of the protective cellular immunity to COVID-19 in different individuals in the population, and points out the importance in designing vaccines to emphasize on the cellular immune response.

Our conclusions are of significance not only for vaccine development, but also for *in vitro* diagnosis. There are increasing reports about variable antibody responses and attenuated humoral immunity in SARS-Cov-2 infected or vaccinated individuals. Memory T cells can remain in the body for a prolonged period of time. Once a vaccine is available, the ELISPOT method may be useful to distinguish between natural infection and vaccination. Furthermore, it could be applied to individuals in the early stage of disease for the evaluation of anti-viral protective immunity, or potentially be useful for risk-stratification in asymptomatic individuals.

## Materials and methods

### Blood donors and PBMC isolation

Healthy donors’ blood specimens were obtained with approval from the Institutional Review Board (IRB) of Shenzhen Geno-Immune Medical Institute (GIMI IRB-20001 and IRB-20002) with informed consent from participants in accordance with regulatory guidelines. PBMCs were isolated using Ficoll-Paque plus (GE Healthcare, Shanghai Co. Ltd).

### Synthesis of overlapping pentadecamer peptides

The eleven overlapping pentadecamer peptides of the SARS-CoV-2 viral proteins were synthesized at >95% purity from Shanghai Royobiotech Co., LTD, which included 47 peptides of the S protein, 53 peptides of the M protein, 16 peptides of the E protein, 38 peptides of the N protein, and 28 peptides of the P protein. The control peptide pool, including viral peptides selected from defined HLA class I-restricted T-cell epitopes for T cell assays, was purchased from JPT Peptide Technologies (Berlin, Germany), and the control HIV-1 Pol peptide pool was gift of NIH AIDS and Reference Reagent Program. The lyophilized peptides were dissolved in DMSO and mixed in equal parts to a final concentration of 1 mg/ml in DMSO before use.

### *In vitro* generation of SARS-CoV-2-specific cytotoxic T lymphocytes

PBMCs were plated into a 10 cm dish at 7×10^7^ cells/dish and adhered for 2 hours in AIM-V (Gibco-BRL, CA). The nonadherent cells were removed gently and frozen as the source of lymphocytes for co-culture use. Adherent monocytes were cultured in AIM-V supplemented with 50 ng/mL of GM-CSF and 25 ng/mL interleukin (IL)-15 (eBiosource International, Inc., Camarillo, CA) for 5 days and incubated for another 24 hours with tumor necrosis factor-α (TNFα, 50 ng/mL), IL-1-β (10 ng/mL), IL-6 (10 ng/mL, all from R&D Systems, MN) and PGE2 (1 mM, Sigma-Aldrich, MO) for maturation. Dendritic cell (DC)-activated antigen-specific immune effector cells were generated as described earlier (36, 37). In brief, mature 2-day DCs were loaded with SARS-CoV-2 S, E, M, N, P pooled peptides (1 mg/mL per peptide) for 3 hours. The antigen-pulsed DCs were co-cultured with autologous nonadherent PBMCs at a ratio of 1:20 in AIM-V with 2% human AB serum. On day 3, half of the medium was replaced with fresh medium supplemented with IL-2 (12.5 U/mL), IL-7 (5 ng/mL) and IL-15 (20 ng/mL, all from Gentaur, Aachen, Germany). Half of the medium was replaced with a fresh medium with cytokines every other day until analysis.

### ELISPOT assay of immune effector function

The T cell precursor frequency against SARS-CoV-2 in health subjects was determined based on the IFN-γ ELISPOT assay. The ELISPOT plate was coated with a capture Ab at 4°C overnight. Then, 1×10^5^ of fresh PBMCs were added to each well of the precoated plate. Next, the specific antigen was added at final 200 μL/well in AIM-V supplemented with 5% hAB without cytokines, at 37°C for 18-24 hrs. The test samples included PBMC or cultured effector cells. PHA activated T cells served as positive control, and HIV-1 pooled peptides were used as control background activity. The concentrations of reagents used were: PHA, final 100ng/mL, SARS-CoV-2 peptides, final 1mg/ml, and HIV-1 pol peptide pool, 1 mg/ml. The cells only without antigen were used as blank and were identified as “Medium”. All samples were tested in duplicate wells. The spots in the plate were quantified using the Bio Reader 400 Pro-X and analyzed in Excel as instructed with standard error. We noted that the E peptides contained many hydrophobic amino acids, which resulted in the addition of trifluoroacetate in the purification of peptides. This made the bottom of ELISPOT plate red. The automated spot reader can distinguish this color background from true spots.

### Immune effector CD107a degranulation and intracellular cytokine staining

These assays were performed as previously described (38). Briefly, 2×10^5^SARS-CoV-2-specific T cells were stimulated for 5 h in a 96 well plate with peptides. Monensin A (Sigma-Aldrich) and FITC-conjugated Abs for CD107a or isotype matched antibodies (BD Pharmingen, San Diego, CA, USA) were added 1 hour after stimulation and incubated for 5 hours. Cells were then stained with antibodies against CD3, CD8, and CD4 and fixed, permeabilized with Cytofix/Cytoperm solution and stained with antibodies against IFN-γ, TNFα and IL-2 (all from BD Pharmingen) at 4°C for 20 min. Unrelated peptide group was included as a negative control for spontaneous CD107a release and/or cytokine production.

### Statistical analysis

Statistical analysis was performed based on student’s t test for two group analysis. P value less than 0.05 is considered statistically significant.

## Author Contributions Statement

LJC designed the studies; CWC, YL, CJ, and HL performed the experiments, CWC drafted the manuscript and LJC, XC discussed the study and made the revisions; all authors read and approved the final manuscript.

## Footnotes

This work is funded by research funds from the the Science and Technology Planning Technical Research Project of Shenzhen (JCYJ20170413154349187, JCYJ20170817172416991 and JCYJ20170817172541842).

